# Generation of synthetic TSPO PET maps from structural MRI images

**DOI:** 10.1101/2024.09.27.615379

**Authors:** Matteo Ferrante, Marianna Inglese, Ludovica Brusaferri, Nicola Toschi, Marco L Loggia

## Abstract

**Background:** Neuroinflammation, a pathophysiological process involved in numerous disorders, is typically imaged using [11C]PBR28 (or TSPO) PET. However, this technique is limited by high costs and ionizing radiation, restricting its widespread clinical use. MRI, a more accessible alternative, is commonly used for structural or functional imaging, but when used using traditional approaches has limited sensitivity to specific molecular processes. This study aims to develop a deep learning model to generate TSPO PET images from structural MRI data collected in human subjects.

**Methods:** A total of 204 scans, from participants with knee osteoarthritis (n = 15 scanned once, 15 scanned twice, 14 scanned three times), back pain (n = 40 scanned twice, 3 scanned three times), and healthy controls (n=28, scanned once), underwent simultaneous 3T MRI and [11C]PBR28 TSPO PET scans. A 3D U-Net model was trained on 80% of these PET-MRI pairs and validated using 5-fold cross-validation. The model’s accuracy in reconstructed PET from MRI only was assessed using various intensity and noise metrics.

**Results:** The model achieved a low voxel-wise mean squared error (0.0033 ± 0.0010) across all folds and a median contrast-to-noise ratio of 0.0640 ± 0.2500 when comparing true to reconstructed PET images. The synthesized PET images accurately replicated the spatial patterns observed in the original PET data. Additionally, the reconstruction accuracy was maintained even after spatial normalization.

**Conclusion:** This study demonstrates that deep learning can accurately synthesize TSPO PET images from conventional, T1-weighted MRI. This approach could enable low-cost, noninvasive neuroinflammation imaging, expanding the clinical applicability of this imaging method.

## 1 Introduction

The translocator protein (TSPO) is a 18 kDa protein primarily expressed on the outer mithochondrial membrane of multiple cell types, and is implicated in multiple physiological and pathological processes [NCA^+^21]. It was initially identified as the “peripheral-type benzodiazepine receptor” [BS77]. However, further studies have revealed that TSPO is extensively distributed throughout various organs in the body, including the brain. In the central nervous system, its expression levels are very low in healthy conditions, but become dramatically upregulated primarily by microglia and/or astrocytes in the context of neuroinflammatory conditions. Because of these expression properties, as well of our ability to image this protein using molecular imaging techniques such as [11C]PBR28 positron emission tomography (PET) imaging, TSPO has been extensively investigated as an in-vivo biomarker for neuroinflammation in various neuropathologies, such as neurodegenerative, psychiatric, chronic pain and other conditions [LCA^+^, ABS^+^22, GRL^+^22, AFS^+^]. However, the clinical utility of TSPO PET is hampered by its high costs, radiation exposure, and infrastructure requirements. In contrast, Magnetic Resonance Imaging (MRI) offers a safer, more accessible alternative. Prior research has shown the capability of MRI to detect some neuroinflammation-associated structural and metabolic changes [AGHL16, Qua15]. However, MRI -in its traditional uses-has low specificity for molecular processes. Leveraging recent advances in deep learning, particularly in medical image translation and synthesis, here we explore the potential of synthesizing TSPO PET images from conventional MRI. Deep learning applications have successfully enhanced and synthesized PET images, as demonstrated in various studies, including conditional Generative Adversarial Networks (GAN) based Computed Tomography (CT) image generation from MRI [WBY^+^], MRI prediction from PET using E-GAN [BRGG22], and synthesizing Florbetapir PET for Alzheimer’s diagnosis [SSVB21]. Moreover, these techniques have facilitated the conversion of low-dose to high-dose PET scans, significantly reducing radiation exposure while maintaining diagnostic accuracy [SJY^+^, SSVB21, ZHQ^+^22].

This study aims to develop a deep learning model for PET image synthesis from T1-weighted MRI, using a 3D U-Net architecture (Figure 1) [CAL^+^]. We choose a 3D U-Net structure over e.g. GANs or diffusion models due to its robust training behavior, ability to capture diverse data distributions, and suitability for limited training datasets. The model’s encoder-decoder structure facilitates the interpretation of imaging features and simplifies the synthesis process, making it a valid choice for PET synthesis from MRI. We therefore propose a tracer-specific modality conversion model based on U-Net, capable of generating synthetic brain PET images from T1-weighted MRI scans. This approach, if successful and validated, would offer a cost-effective, non-invasive method for imaging neuroinflammation, providing new insights into MRI features relevant to glial activation and related processes. Our work demonstrates the feasibility of synthesizing TSPO PET images from structural MRI images, hence significantly contributing to the non-invasive characterization of neuroinflammation.

**Figure 1:**
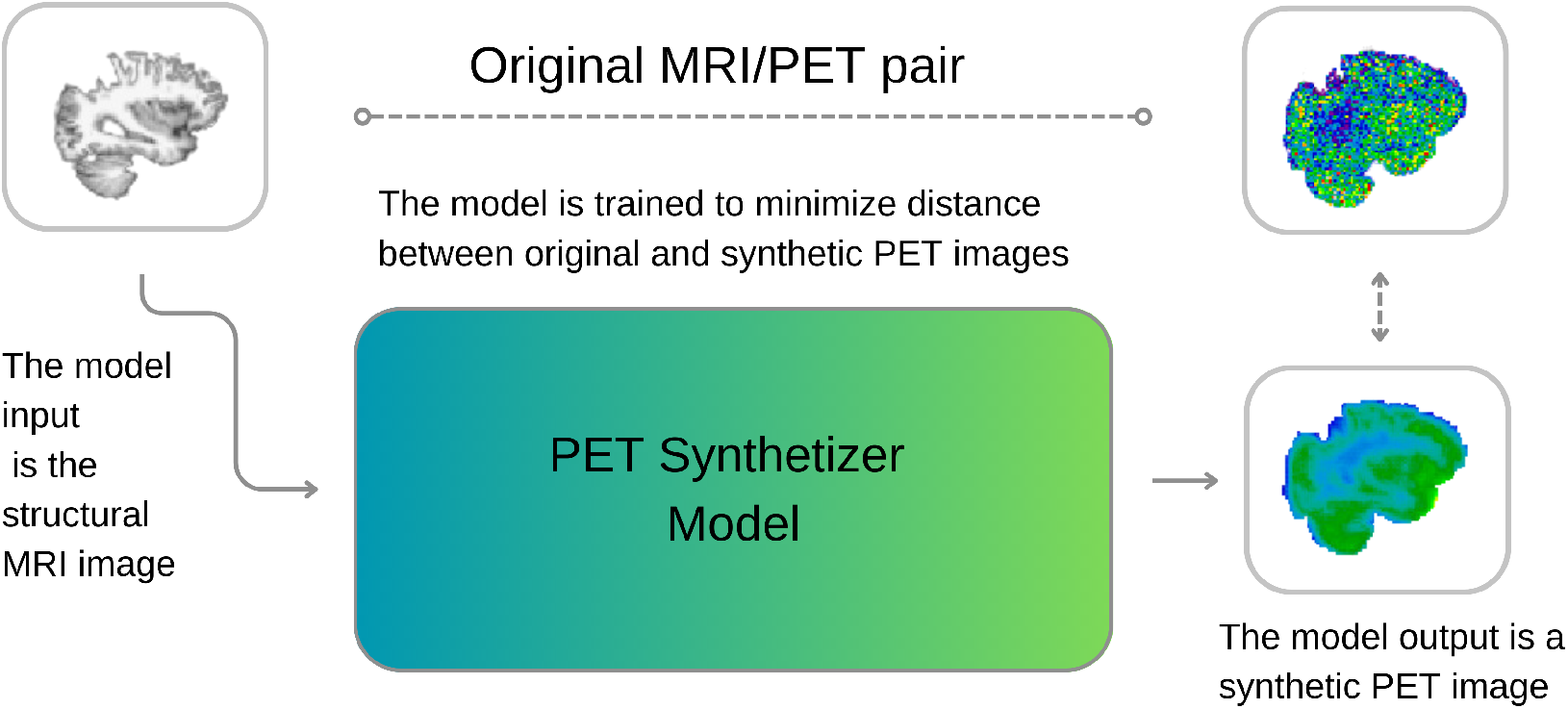
A visual description of our approach. During training, the model learns to synthesize PET images from MRI, minimizing the discrepancies between original and reconstructed images. During inference, the model can generate new synthetic PET images from structural T1w MRI images.

## 2 Material and Methods

This section outlines our experimental setup. The code, developed in Python 3.9, leverages deep learning libraries such as Pytorch and Monai [CLB^+^22]. All computations were conducted on a server with four NVIDIA A100 GPUs (80GB RAM each) and 2 TB of System RAM.

### 2.1 Dataset

A total of 204 scans, from participants with knee osteoarthritis (n=15 scanned once, 15 scanned twice, 14 scanned three times), back pain (n = 40 scanned twice, 3 scanned three times), and healthy controls (n=28, scanned once) were included for this study. They underwent simultaneous 3T MRI and TSPO PET neuroimaging with [11C]PBR28. During cross validations (see below) when an individual had multiple scans, care was taken to segregate all scans from that individual either in the training or in the test split. Each participant received 9-15 mCi of [11C]PBR28 intravenously, and was scanned using a Siemens Biograph mMR for 90 minutes. From each scan, a standardized uptake value (SUV) map was reconstructed using 60-90 minute post-injection data, which were attenuation-corrected using a T1-weighted structural data (multi-echo magnetization-prepared rapid acquisition with gradient echo (MPRAGE); TR/TE1/TE2/TE3/TE4 = 2530/1.64/3.5/5.36/7.22ms, flip angle = 7°, voxel size = 1mm isotropic) and the PseudoCT approach [IGHF^+^14].

### 2.2 Pre- and Post-processing

Our preprocessing pipeline, including alignment, skull stripping, and normalization, was performed using FSL [JBB^+^12]. Coregistration aligned both modalities to the same native space. Skull stripping, performed next, isolated the brain to eliminate non-brain signals. MRI volumes were normalized to a [0,1] scale using robust scaling (1st and 99th percentiles) via MONAI, while PET volumes were normalized to SUV units based on individual minimum and maximum intensities. This approach harmonized intensities across subjects, thereby facilitating convergence in model training. Additionally, genotype data (high-affinity and mixed-affinity binder status, which affects [11C]PBR28 binding was used to adjust PET binding variability using FSL by regressing out the effect of genotype before further analysis [[OYG^+^12]). We implemented a 5-fold cross-validation strategy, dividing the 204 cases into five subsets. Each fold uses 80% of the data for training and 20% for testing, rotating across all subsets. The model’s objective (see “Model” section) is to synthesize TSPO SUV maps from structural MRI, while the reconstruction performance is evaluated against real PET images in the test set (see “model Evaluation” section). This validation approach ensures robust assessment of the model’s capability to generalize from MRI to PET in our diverse patient cohort. To ensure a comprehensive and robust assessment of the synthesized PET image quality and the efficacy of our deep learning model (see below), a linear registration to MNI space and a nonlinear warping were applied to all T1 MRI images together with the true and synthesized PET images. This consistent registration enabled a thorough and standardized evaluation of the synthesized PET images against the true PET targets, using intensity, asymmetry, noise, and regional metrics as detailed below. Furthermore, under the hypothesis that the reconstruction process could filter out unstructured noise, we also compared reconstructed PET images to smoothed real PET images (4 mm FWHM kernel Gaussian filter).

### 2.3 Model

The 3D U-Net architecture was specifically adapted for the task of synthesizing positron emission tomography (PET) images from magnetic resonance imaging (MRI) inputs. The core of the model’s design is a depthwise separable convolution-based encoder, which incorporates four downsampling layers. This approach minimizes the number of parameters in the model, thereby enhancing computational efficiency and facilitating the handling of three-dimensional data while reducing risk of overfitting. Each layer in the encoder and decoder pathways is comprised of 32 channels, optimizing the model’s capacity to capture a wide range of features in 3D medical images while conserving RAM usage.

Depthwise separable convolutions, a key component in our model, partition the standard convolution operation into two stages: depthwise convolutions and pointwise convolutions. This strategy, as discussed by [Cho17], drastically lowers the parameter count, thereby allowing for the inclusion of more channels in each layer without a proportional increase in computational demand. This technique is therefore instrumental in achieving high efficiency in feature extraction from volumetric data.

Moreover, the final layer of the encoder integrates a self-attention mechanism, thereby augmenting the model’s capability to understand global contextual relationships alongside local feature extraction. This enhancement is key for capturing long-range dependencies across the volumetric space of the input data, and is hypothesized to aid in capturing underlying biological and anatomical structures.

On the decoder side, the architecture uses transpose convolutions for the purpose of upsampling, effectively restoring the spatial resolution that is reduced during downsampling. Skip connections, bridging corresponding layers in the encoder and decoder, are employed to reintroduce high-resolution details and features into the reconstructed images, ensuring that the latter can exploit the intricate details present in the original MRI scans.

The training of the model was conducted over 50 epochs, using the Adam optimizer, chosen for its well documented efficiency in converging on complex optimization landscapes. The loss function we employed is a hybrid formulation that combines binary cross-entropy (BCE) and mean squared error (MSE), defined as follows:

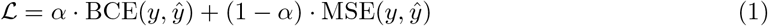

where *y* denotes the true PET images, *ŷ* represents the synthesized PET images, and *α* is a weighting factor that balances the contribution of each component to the total loss. This dual loss function is designed to ensure fidelity both in terms of the visual similarity and the voxel intensity distributions between the synthesized and true PET images, thus addressing both qualitative and quantitative aspects of the image synthesis task.

### 2.4 Model Evaluation

To rigorously assess our model’s performance in PET image synthesis, we employed a comprehensive framework comprised of various quantitative metrics, which are computed both at voxel level and at parcellation level. The latter, region-of-interest (ROI)-wise analysis was conducted using the cortical and subcortical Harvard-Oxford atlases [DSF^+^06].

Voxel-wise analysis incorporated the mean squared error (MSE) to gauge reconstruction accuracy, with percentage difference maps highlighting spatial error distribution. We also employed a metric called normalized difference (NormDiff):

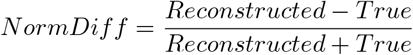

This metric evaluates relative errors, disregarding absolute intensity levels and has the advantage of being bounded between −1 and 1.

Additionally, we included contrast-to-noise ratio (CNR) was calculated due to its clinical relevance in diagnostic imaging. In this paper, the following definition of CNR was adopted:

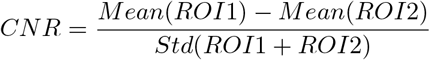

Since CNR varies locally, to produce a reliable global estimate for each image pair we proceeded as follows: we sampled ROI1 and ROI2 times (size: 4×4×3 voxels) at random 1000 times in each PET image, computed CNR for each ROI, and compared the median CNR between the original and the reconstructed image.

## 3 Results

Table 3 and Figure 2 present a detailed comparison of synthesized PET images against true PET targets. Figure 3 shows the results of the ROI-wise analysis in terms of normalized difference, MSE, percentage difference analysis and contrast-to-noise ratio evaluation. We achieved a low MSE of 0.0034 ± 0.0010 (to be compared with the input and output values which span the interval [0,1]) for raw synthesized PET, indicating minimal voxel-wise intensity error Smoothing the synthesized PET images further reduces MSE.

**Figure 2:**
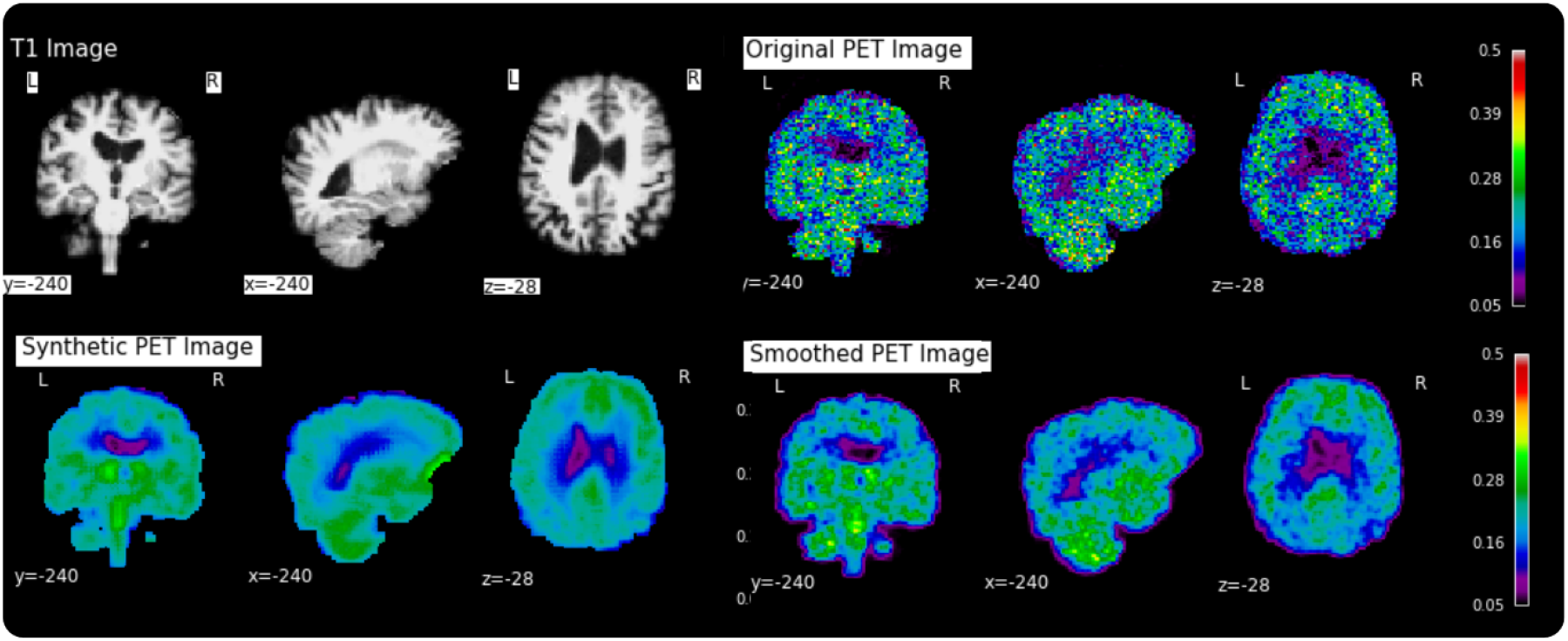
Examples of a random subject from the test set. The first row displays original intensity-normalized structural MRI and PET images, the second row presents our reconstructed images (left) and the original PET image post Gaussian smoothing (fwmh = 4mm), (right).

**Figure 3:**
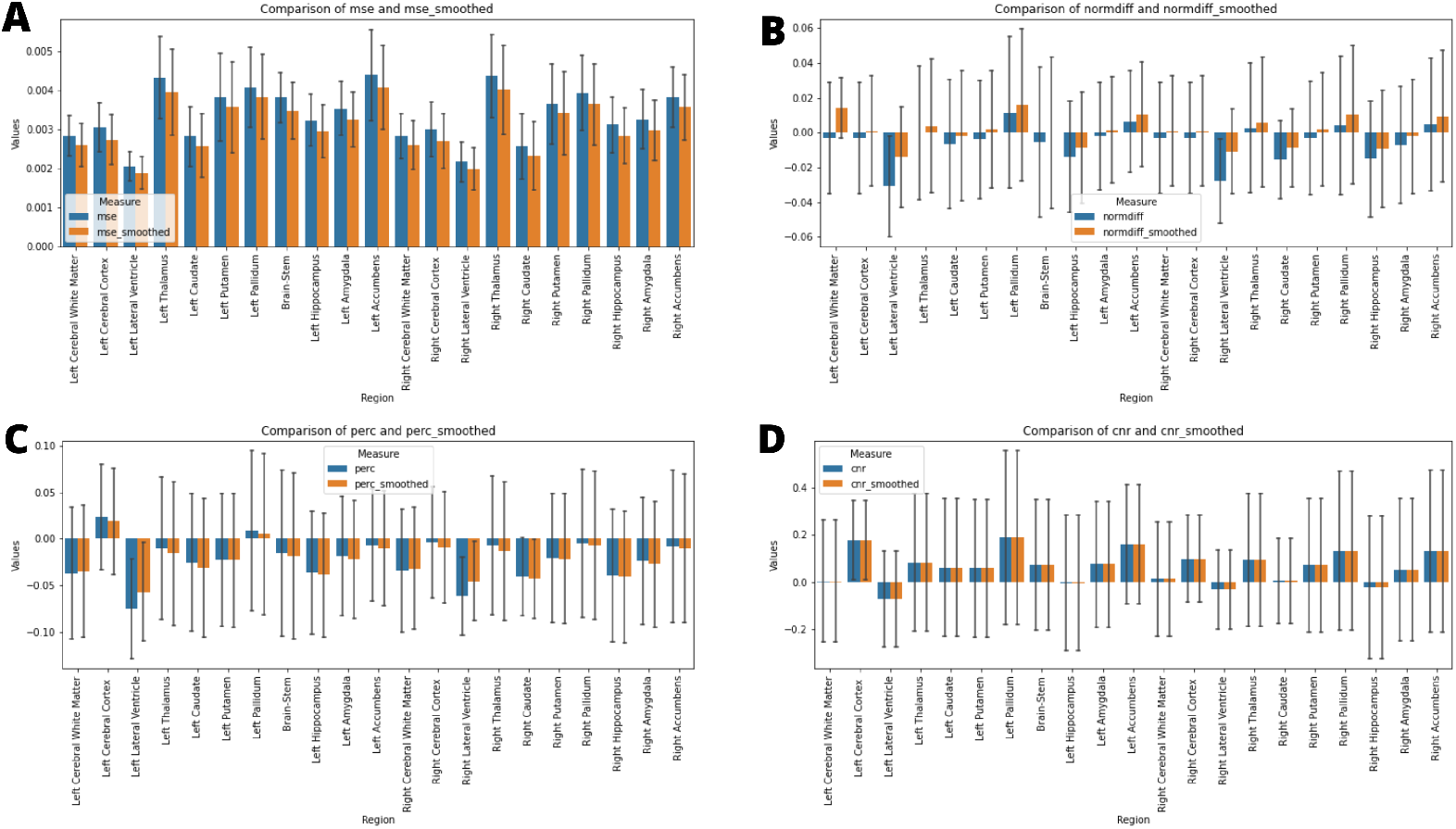
Results of ROI-wise analysis, which compared original to reconstructed PET images. The error bars report standard error across folds. **A**: MSE comparison between synthetic images and original/smoothed. **B**: Normalized Difference (NormDiff) between original and smoothed images. **C**: Percentage difference analysis between original and smoothed images. **D**: Median CNR comparison between original and smoothed images.

The normalized Difference (NormDiff), showed a mean close to zero, confirming an unbiased reconstruction of PET SUV maps. Smoothing the synthesized images further lowered this measure, suggesting that our model could potentially also function as a signal enhancing smoothing filter.

CNR calculations yielded similar mean values across raw and smoothed synthesized images, demon-strating the preservation of spatial characteristics in synthesized PET.

The mean percentage difference between synthesized and true PET was modest, with a slightly more negative bias observed after applying the smoothing filter. These results collectively underscore the model’s ability to synthesize PET images with high accuracy and minimal bias, reflected in low global error magnitudes and preserved intensity characteristics. The efficacy of the model is further evidenced by the consistent quality of PET synthesis across subsets, even after spatial normalization to MNI space, highlighting its robustness.

**Table 1:**
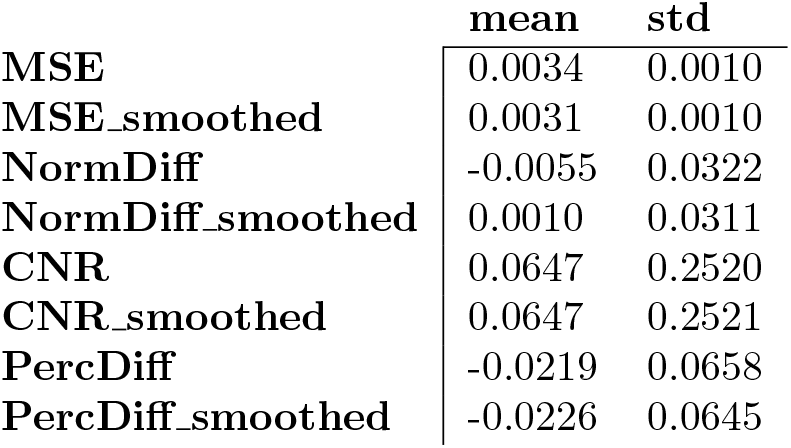
Quantitative metrics (MSE, NormDiff, CNR, and percentage difference (PercDiff)) comparing synthesized PET to true PET images across the test set. Results include both raw synthesized PET outputs and a smoothed version using a Gaussian filter with a 4 mm kernel.

|  | mean | std |
| --- | --- | --- |
| MSE | 0.0034 | 0.0010 |
| MSE_smoothed | 0.0031 | 0.0010 |
| NormDiff | -0.0055 | 0.0322 |
| NormDiff_smoothed | 0.0010 | 0.0311 |
| CNR | 0.0647 | 0.2520 |
| CNR_smoothed | 0.0647 | 0.2521 |
| PercDiff | -0.0219 | 0.0658 |
| PercDiff_smoothed | -0.0226 | 0.0645 |

## 4 Discussion

Neuroinflammation is being increasingly recognized for its role in a multitude of brain conditions; as such, monitoring this process may prove valuable in many clinical contexts. However, PET-based imaging of neuroinflammation -currently the best tool at our disposal-is hindered by high costs, and the need to expose patients to ionizing radiation. In contrast, MRI, which is safer and more widely available, could in principle offer an accessible alternative with its capability to detect neuroinflammation signal. We developed a deep learning model for PET image synthesis from T1-weighted MRI to transform widely available MRI scans into informative PET-like neuroinflammation patterns. We adopted a 3D U-Net architecture for its stable training performance, its efficiency with small datasets and ability to capture diverse data distributions. In addition, the encoder-decoder design of the U-Net model allows an easy interpretation of the extracted imaging features simplifying the synthesis process. We obtained a low voxel-wise mean squared error (0.0033 ± 0.0010) across all folds together with a median contrast-to-noise ratio of 0.0640 ± 0.2500.

The results of this study have important implications for neuroinflammation imaging and chronic pain management. We introduced a novel and non-invasive method that could transform current diagnostic and therapeutic strategies for chronic pain conditions offering an opportunity to monitor neuroinflammation in chronic pain patients more frequently with a reduced cost. This could not only facilitate personalized treatment but also the acquisition of large-scale studies, potentially leading to a better understanding of the complex mechanisms of chronic pain and to the discovery of new therapeutic targets. Our model estimates spatial patterns of TSPO PET signal which align with existing research indicating variable glial activation in different pain pathologies [LCA^+^], suggesting MRI’s promise in capturing such variations.

In this study, we used the SUV as the standardized measure of PET tracer uptake. However, the debate on the optimal PET quantification methods remains ongoing [Key95] and different metrics are frequently employed across studies. For example, the SUV ratio (SUVr) could be explored in future applications to allow for a more precise analysis of tracer binding in the brain by comparing it with a reference region [YTH^+^15]. Additionally, moving beyond TSPO tracers to newer markers like P2X7, COX-2 could significantly enhance our understanding of microglial activation, paving the way for a multi-tracer PET synthesis approach [YFD^+^06] which would significantly enhance the availability of neuroinflammation assessment across centers. Furthermore, the integration of multi-parametric MRI could provide complementary information to further enhance the synthesis of PET images by including unique pathological features into the reconstruction process. Future research will also focus on the validation of synthesized TSPO PET images against clinical outcomes to validate the clinical utility of the synthesized images in guiding treatment decisions. Additionally, exploring the model’s applicability to other neuroinflammatory conditions beyond chronic pain could broaden its impact, making it a versatile tool in neurology and psychiatry.

## 5 Conclusion

This study illustrates the successful synthesis of TSPO PET SUV images from structural MRI in chronic pain patients and healthy volunteers, marking a significant advance in non-invasive neuroimaging. The synthetic PET images generated by our model reproduce spatial signal distributions and related contrast properties which are extremely close to real PET scans.

In addition, synthesized PET volumes, while smoother than the original data, closely resembled Gaussian-smoothed PET scans. Given the conventional PET processing commonly includes smoothing, it is possible to hypothesize that our model inherently filters out unstructured noise, potentially enhancing signal-to-noise ratio and thereby the sensitivity for detecting subtle neuroinflammatory differences.

In essence, our findings demonstrate that deep learning can effectively transform widely available MRI scans into informative PET-like neuroinflammation patterns. This approach holds promise for improving the noninvasive study of glial involvement in chronic pain and potentially other conditions, offering a novel perspective in medical imaging research.

## 6 Acknowledgements

This work is supported and funded by the National Institutes of Health (NIH) through grants R01NS095937 (to MLL), R01DA053316 (to MLL) and R01NS094306 (to MLL); NEXTGENERATIONEU (NGEU); the Ministry of University and Research (MUR); the National Recovery and Resilience Plan (NRRP); project MNESYS (PE0000006, to NT) - A Multiscale integrated approach to the study of the nervous system in health and disease (DN. 1553 11.10.2022); the MUR-PNRR M4C2I1.3 PE6 project PE00000019 Heal Italia (to NT); the NATIONAL CENTRE FOR HPC, BIG DATA AND QUANTUM COMPUTING, within the spoke “Multiscale Modeling and Engineering Applications” (to NT); the European Innovation Council (Project CROSSBRAIN - Grant Agreement 101070908, Project BRAINSTORM - Grant Agreement 101099355); the Horizon 2020 research and innovation Programme (Project EXPERIENCE - Grant Agreement 101017727). Matteo Ferrante is a Ph.D. student enrolled in the National PhD in Artificial Intelligence, XXXVII cycle, course on Health and Life Sciences, organized by Universita Campus Bio-Medico di Roma.

